# A comprehensive analysis of humanized mouse models for the study of cancer immunotherapies

**DOI:** 10.1101/2025.09.04.674233

**Authors:** Philippe De La Rochere, Laure Loumagne, Melanie Rathaux, Marine Dubois, Fariba Nemati, Sophie Viel, Tamara Slavnic, Jayant Thatte, Qixiang Li Henry, Xuesong Ouyang, Christine Sedlik, Didier Decaudin, Georges Azar, Sukhvinder Sidhu, Eliane Piaggio

**Author notes:** Equal author contribution (Philippe De La Rochere, Laure Loumagne). genOway, 31 rue Saint-Jean-de-Dieu, Lyon F-69007, France (Philippe De La Rochere). Institut Servier d’Innovation Thérapeutique, 22 route 128, Gif-sur Yvette, F-91190, France (Melanie Rathaux). Egle Therapeutics, 5 Rue Gardenat Lapostol, Suresnes F-92150, France (Marine Dubois). Plateforme CYM, IPSIT, Université Paris Saclay, 17 avenue des Sciences, Orsay F-91400, France. **Correspondence:** Eliane Piaggio INSERM U932, Institut Curie, 26, rue d’Ulm, 75005 Paris, France.

## Abstract

Humanized immune system (HIS) mouse models, generated by engrafting tumors and hematopoietic cells of human (Hu) origin into immunodeficient host mice, effectively recapitulate key aspects of the crosstalk between human immune cells and tumors. These models represent a valuable tool for the preclinical evaluation of immunotherapies. In this study, we provide a comprehensive comparison of two widely used HIS models: the Hu-CD34+ model, which engrafts Hu-hematopoietic cells derived from Hu-CD34+ hematopoietic stem cells (HSCs), and the Hu-PBMC model, which utilizes Hu-peripheral blood mononuclear cells (PBMCs). We assess the kinetics, quality and extent of immune cell engraftment, as well as the development of graft-versus-host disease (GVHD). Additionally, we investigate the impact of different immunodeficient host mouse strains on immune cell reconstitution in the Hu-CD34+ model. Both HIS models were engrafted with human tumors derived from either cell lines or patient-derived xenografts (PDX), revealing distinct immune-tumor interactions that influenced antitumor responses. Notably, tumor responses to T-cell-directed therapies, including anti-PD1 antibodies, IL-2-anti-IL-2 antibody complexes, and T-cell engagers, varied across these models. Our findings provide novel insights into the properties and limitations of HIS models, offering a critical resource for optimizing next-generation immuno-oncology strategies and guiding the design of future therapeutic interventions.

## Introduction

The blockade of immune checkpoints with antibodies (anti-CTLA-4, anti-PD1, anti-PD-L1) has shown impressive clinical outcomes, establishing immunotherapy as a promising cancer treatment (1,2). Yet, progress is hindered by the lack of translationally relevant in vitro and in vivo models. Developing mouse tumor models is critical to study mechanisms of action, identify biomarkers, and prioritize combinatorial approaches with targeted therapies, radiotherapy, or chemotherapy.

Current preclinical models include syngeneic tumors and genetically engineered mice in immune-competent hosts, and humanized immune system (HIS) models in immunodeficient mice (3). While the first two have been widely used, they depend on the murine immune system, which does not fully recapitulate its human immunity and limits testing of non-cross-reactive agents (4). HIS models, which reconstitute a functional human immune system and support human tumor engraftment, provide a more physiologically relevant platform (5,6).

HIS models are typically generated with CD34+ hematopoietic stem cells (Hu-CD34+) or peripheral blood mononuclear cells (Hu-PBMC). CD34+ cells enable long-term reconstitution of multiple immune subtypes, though functionality remains incomplete. Improved strains (NSG, NSG-SGM3, NOG-EXL, BRGS, BRGSF) have been developed, but systematic comparisons remain limited (7–12). In contrast, the Hu-PBMC model mainly reconstitutes T cells but is restricted by acute xeno-graft-versus-host-disease (GvHD)within 3-5 weeks (13).

Key hurdles include persistence of murine innate immune cells, difficulties in obtaining matched donor–tumor samples, incomplete reconstitution, GvHD, and high cost. Nonetheless, HIS mice uniquely allow assessment of human immune responses, patient heterogeneity, and personalized immunotherapies.

In this study, we compared CD34+ HSC- and PBMC-based HIS models across mouse strains, focusing on immune reconstitution, tumor interactions, and immunotherapy responses. Our findings provide guidance for their application in immuno-oncology and personalized cancer therapy.

## Materials and Methods

### Mice

NOD.Cg-*Prkdc^scid^ Il2rg^tm1Wjl^*/SzJ (NSG) and NOD.Cg-*Prkdc^scid^ Il2rg^tm1Wjl^* Tg(CMV-IL3,CSF2,KITLG)1Eav/MloySzJ (NSG-SGM3) (Jackson Laboratory) and NOD.Cg-*Prkdc^scid^ Il2rg^tm1Sug^*/JicTac (NOG) and NOD.Cg-*Prkdc^scid^ Il2rg^tm1Sug^* Tg(SV40/HTLV-IL3,CSF2)10-7Jic/JicTac (NOG-EXL) (Taconic Biosciences) were used. Procedures were approved by the ethics committees of SANOFI CEPAL-CRVA #21 (APAFiS #5644), and of Institut Curie CEEA-IC #118 (APAFiS #15227 and APAFiS#25870) following UKCCCR and international guidelines or conducted at Crown Bioscience (protocol #EB17-030-002) approved by the institutional animal care and use committee (IACUC).

### Human immune cells

Hu-PBMCs were isolated from buffy coats (Etablissement Français du Sang), Hu-CD34+ cells from cord blood (ABCell-Bio, France). Human samples followed the Declaration of Helsinki and national regulations.

### Hu-CD34^+^ HSC-mice

Mice were sub-lethally irradiated: 1.4Gy for 3-week-old NSG, 1.1Gy for 3-4-week-old NOG, 1 Gy for 5-week-old NSG-SGM3, and 0.6 Gy for 4-6-week-old NOG mice. Eighteen to 24 hours post-irradiation, mice were injected intravenously (iv) with human CD34^+^ HSC cells (1.3×10^5^ cells for NSG and NOG, 1×10^5^ for NSG-SGM3, and 0.5×10^5^ for NOG-EXL mice). In some experiments (Figure 3), CD34⁺ HSC–humanized NSG mice were obtained from Jackson Laboratory.

### Hu-PBMC-NSG mice

NSG mice were grafted with human tumor cell lines (CDX) or patient-derived xenografts (PDX) and then iv injected with Hu-PBMCs from 2 independent donors in separate mice. Hu-PBMCs were isolated using Ficoll gradient (lymphoprep, Stemcell).

### Tumor cell lines and patient-derived xenografts

Human triple negative breast cancer MDA-MB231 cell line, NSCLC cancer HCC827 cell line (ATCC) and HT-29 colorectal cell line (ATCC) were grown in RPMI medium (Life Technologies) supplemented with 10% heat-inactivated fetal bovine serum (Biosera), glutamine and penicillin-streptomycin (Life Technologies). 3×10^6^ HT-29 or 5×10^6^ MDA-MB231 or HCC827 tumor cells were injected subcutaneously (sc) and tumor growth was measured using a caliper twice weekly.

PDXs were obtained following informed consent from patients undergoing surgery. PDX fragments were grafted into the interscapular fat pad, except in Fig. 3 where tumors from PDX were dissociated using Collagenase / DNase into single cell suspension and 100-200×10^3^ cells were mixed with Cultrex matrix at 1:1 ratio and implanted sc. NSCLC and breast PDX models are available via EuroPDX and PDXFinder, SCLC PDX from Crown Bioscience.

### *In vivo* treatments

Anti-PD1 mAbs (Nivolumab (Opdivo^®^) or Pembrolizumab (Keytruda^®^) were injected intra-peritoneally (ip), biweekly, starting 3-10 days post-PBMC injection, at 10mg/Kg. Control mice were untreated or received human IgG4 isotype antibody. IL-2Cxs (14) were injected ip for five days, 5 days after PBMCs, then biweekly. EpCAM-CD3 Ab was injected iv at 2.5mg/kg biweekly.

### Flow cytometry

Tumor, spleen, bone marrow (BM), and PB samples were processed, red blood cells-lysed, and stained. Tumors were digested with DNAse (10mg/ml, Roche), and liberase LT (5mg/ml, Roche), dissociated (Gentlemacs, Miltenyi).

Cells were incubated with human and mouse Fcγ-receptor blockers (Human TruStain FcX™ (Biolegen), rat anti-mouse CD16/32 (BD Biosciences) and stained for surface markers using: huCD4-BUV395 clone SK3, huCD8-BUV496 clone RPA-T8, huCD45-BUV805 clone HI30, huCD56-BV421 clone NCAM16.2, huCD3 BV510 clone UCHT1, huCD19-BV650 clone SJ25C1, huCD16-FITC clone 3G8, huCD33-PECF594 clone WM53, mCD45-PECy7 clone 30-F11, huCD14-AF700 clone M5E2, huCD45RA-BV421 clone HI100, huCD197-PECF594 clone 150503, huCD45RO-APC clone UCHL1, huCD8-PECF594 clone RPA-T8, huCD56-PE-Cy5 clone N901, huCD45-APC Cy7 clone 2D1, huCD19-Alexa 700 clone HIB19, huTCRgd-FITC clone 11F2 purchased from BD Biosciences; and uhCD25-PE clone M-A251, huCD27-BV605 clone O323, huCD3-BV650 clone OKT3, huCD4-BV785 clone OKT4, huPD-1-BV711 clone EH12.2H7, and huCD25-BV786 clone BC96 from Biolegend; huHLA-DR-FITC clone AC122 from Miltenyi Biotec and huCD45RA-PECy5 clone HI100 from eBiosciences.

Dead cells were stained with LIVE/DEAD Fixable Aqua (Life TechnologiesTM) or Viability Dye eFluor 780 (eBiosciences). Fortessa (BD Biosciences) was used for cell acquisition and FlowJo software v10.0.8 (TreeStar) for data analysis.

### Plasma cytokine measurement

EDTA-plasma was analyzed using the MSD U-PLEX assay platform (U-PLEX Proinflam Combo 1 Human K15049K; Mesoscale Discovery). Plates were analyzed in MSD Discovery Workbench.

### Statistical analyses

For tumor volume data (except Fig 3A), a Two-Way Analysis of Variance Type (ANOVA Type) was performed on the changes from baseline when baseline was available, and on raw tumor volumes otherwise with time as a repeated factor. For Fig 3A, a mixed effect model with factor group, day (continuous) and lung tumor model, as well as all two-way interactions, was performed on log-transformed tumor volumes. Percentage of cells and body weights were analyzed with a Two-Way Analysis of Variance (ANOVA) on raw or transformed data depending on their distribution with time as a repeated factor (except for Fig 6E). For cytokine concentration and % cells measured at week 18 only, a One-Way ANOVA was performed on log-transformed data with time as a repeated factor for cytokines and with a random effect on donors for % cells. For all models, adjustment for multiplicity was applied when multiple comparisons were performed.

Analyses were performed using SAS® version 9.4 for Windows 7 and graphs were generated with GraphPad Prism v7.

## Results

### 1- Characterization of the Hu-CD34^+^ model: immune cell reconstitution and GvHD development

Few studies have directly compared immune cell reconstitution across strains (Maser 2020). We conducted a side-by-side comparison of the four most used genetically modified immunodeficient mice: NSG, NSG-SGM3, NOG, and NOG-EXL, which differ in their ability to support human immune cell development, particularly in the myeloid versus lymphoid compartments. The NSG and NOG mice differ in their genetic modifications, but both achieve a functional knockdown of the IL2R gamma chain, resulting in a profound immunodeficiency with absent T, B, and NK cells. The NSG-SGM3 and NOG-EXL strains are transgenic for the expression of human IL-3 and GM-CSF for both strains and KIT-ligand for NSG-SGM3 only, which enhances the differentiation and survival of human myeloid cells.

All strains showed stable engraftment of Hu-CD45+ cells in blood up to 18 weeks post-HSC injection (Fig. 1 and Fig. S1A-E). Although the kinetics of Hu-CD45+ reconstitution was similar across strains, NSG-SGM3 mice exhibited faster reconstitution early on, though by week 18 all strains reached 50–70% Hu-CD45+.

**Figure 1:**
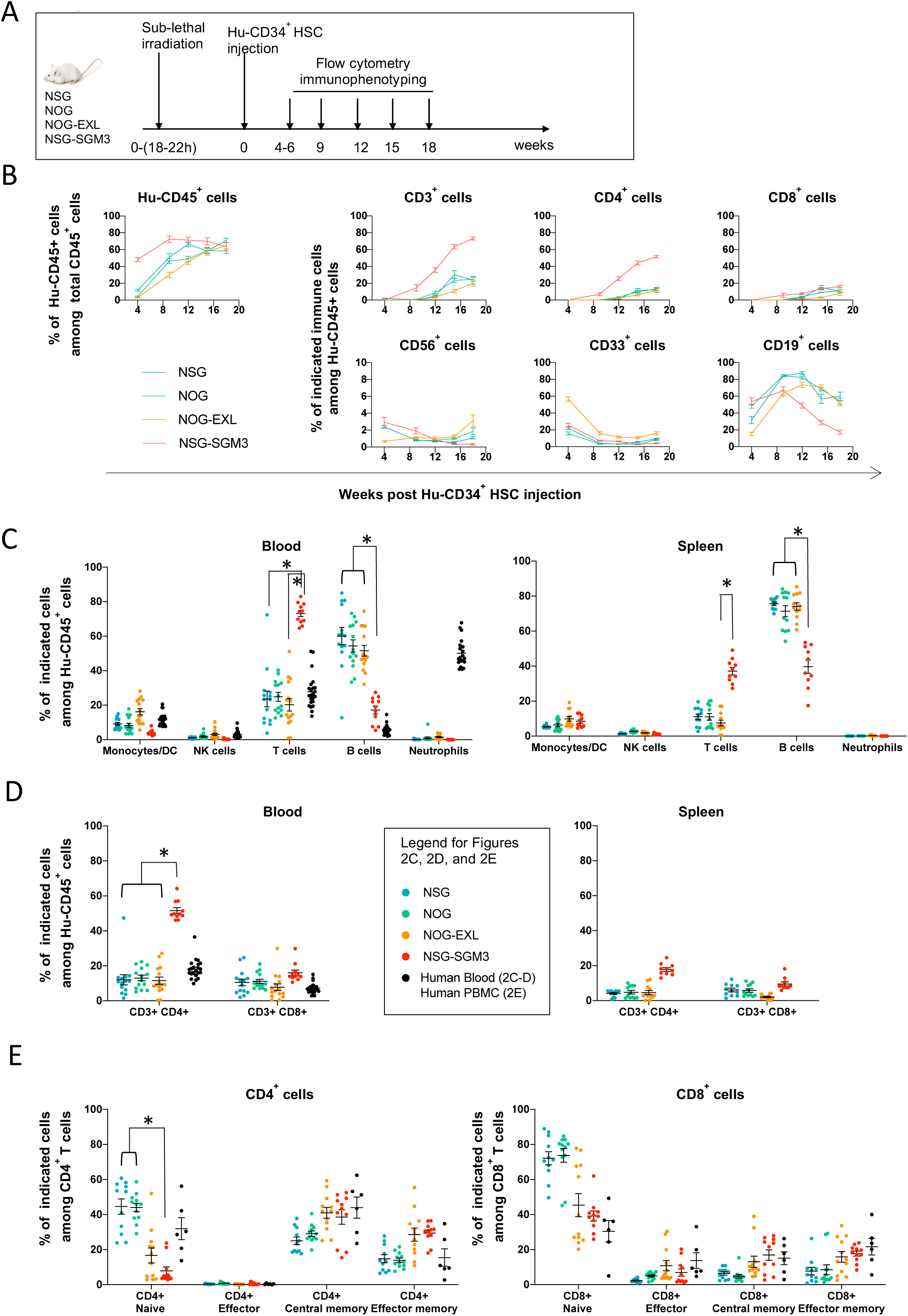
Human immune cell reconstitution upon Hu-CD34^+^ HSC injection in different strains of immunodeficient mice. **A.** Experimental design. 50-130×10^3^ Hu-CD34^+^ HSC were injected into NSG, NOG, NOG-EXL or NSG-SGM3 mice. Blood was recovered at weeks 4 to 6, 9, 12, 15 and 18 post-injection of Hu-CD34^+^ HSC to quantify human reconstitution by flow cytometry analysis. **B.** Frequencies (%) of Hu-CD45^+^ cells population relative to total CD45^+^ cells (Hu-CD45^+^ + m-CD45^+^) and subpopulations among Hu-CD45^+^ cells, in blood over time. **C, D.** Distribution of the Hu-CD45^+^ cell subpopulations relative to Hu-CD45^+^ cells in blood and spleen of different mouse strains. Mice were sacrificed 18 weeks after Hu-CD34^+^ HSC injection and the distribution of the immune cell populations was determined by flow cytometry. Monocytes/DC are defined as Non neutrophils CD33^+^ cells, NK cells as CD3^-^ CD56^+^ cells, T cells as CD3^+^ CD56^-^ cells, B cells as CD19^+^ cells and Neutrophils as SSC^high^ CD16^high^ cells as described in the gating strategy (Supplementary Fig. S1A). **E.** T cell differentiation state in Hu-CD34^+^-humanized mice. Frequencies (%) of naïve, memory, effector memory, and effector cells relative to CD4^+^ T cells (left panel) and CD8^+^ T cells (right panel) were determined by flow cytometry analysis. Gating strategy is similar to the one used for Hu-PBMC-NSG mouse as illustrated in supplementary Figure 2C. Data were obtained from mice reconstituted with 2 or 3 different Hu-CD34+ HSC donors with n=10-15 mice per mouse strain for blood and spleen data. Data is expressed as individual dots and mean ± SD. *p<0,05 was obtained with a One-Way Analysis of Variance on log-transformed with Tukey’s correction for multiplicity.

Immune cell subsets displayed different engraftment kinetics. NK (CD56+), non-neutrophil myeloid cells (CD33+) and B cells (CD19+) were detectable as early as week 4 post-humanization, whereas T cells (CD3+) were not observed until week 12 in most strains, except for NSG-SGM3, which showed earlier and higher T-cell reconstitution, particularly CD4+ T cells (**Fig. 1B**). Notably, in NSG-SGM3 mice, B-cell reconstitution was significantly lower than in the other strains. Furthermore, 7 out of 19 NSG-SGM3 humanized mice (39%) requiered euthanasia due to severe weight loss between weeks 10 and 17 post-humanization (data not shown), suggesting toxicity.

At week 18 post-HSC injection, immune cell distribution in blood and spleen was assessed relative to human donor blood. T, NK, and non-neutrophil myeloid cells (monocytes/DC) were present in similar proportions to those in healthy donors, while B cells were overrepresented in all strains except NSG-SGM3 (**Fig. 1C**). Neutrophils remained absent in circulation across all models, and despite enhancements in myeloid differentiation in NSG-SGM3 and NOG-EXL strains, no significant improvements in myeloid reconstitution were observed at this time point (**Fig. 1C and Fig. S1E-G**). Human neutrophils were detected in bone marrow of all strains, albeit at significantly lower frequencies in NSG-SGM3 mice (**Fig. S1F**).

Analysis of T-cell subsets revealed that NSG-SGM3 mice had a significantly higher proportion of CD4+ T cells, and a reduced proportion of B cells (**Fig. 1D**). The overall T-cell compartment in all models exhibited normal frequencies of naïve, effector, central memory, and effector memory cells, although NSG-SGM3 mice displayed a relative depletion of naïve T cells (**Fig. 1E**). In contrast to human donor blood, NK cells in all models lacked the characteristic predominance of cytotoxic NK cells over cytokine-producing NK cells (**Supplementary Fig. S1B-C**).

Importantly, no NSG, NOG, or NOG-EXL mice displayed clinical signs of GvHD throughout the study period, supporting their use for long-term immunotherapy evaluation.

### 2- Characterization of the Hu-PBMC model: immune cell reconstitution and GvHD development

The Hu-PBMC-NSG model is widely used to study human immune responses, but limited by inevitable GvHD, caused by human T cells attacking murine tissues(15,16). Therefore, we sought to define an optimal therapeutic window between immune reconstitution and overt GvHD onset.

To optimize PBMC dosing for delayed GvHD onset, we injected non-irradiated NSG mice with varying PBMC doses and monitored immune engraftment and GvHD (**Fig. 2A and Fig. S2A**). Stable reconstitution was defined as ≥2% Hu-CD45+ cells at two time points. GvHD onset was defined when weight loss was >20%.

**Figure 2:**
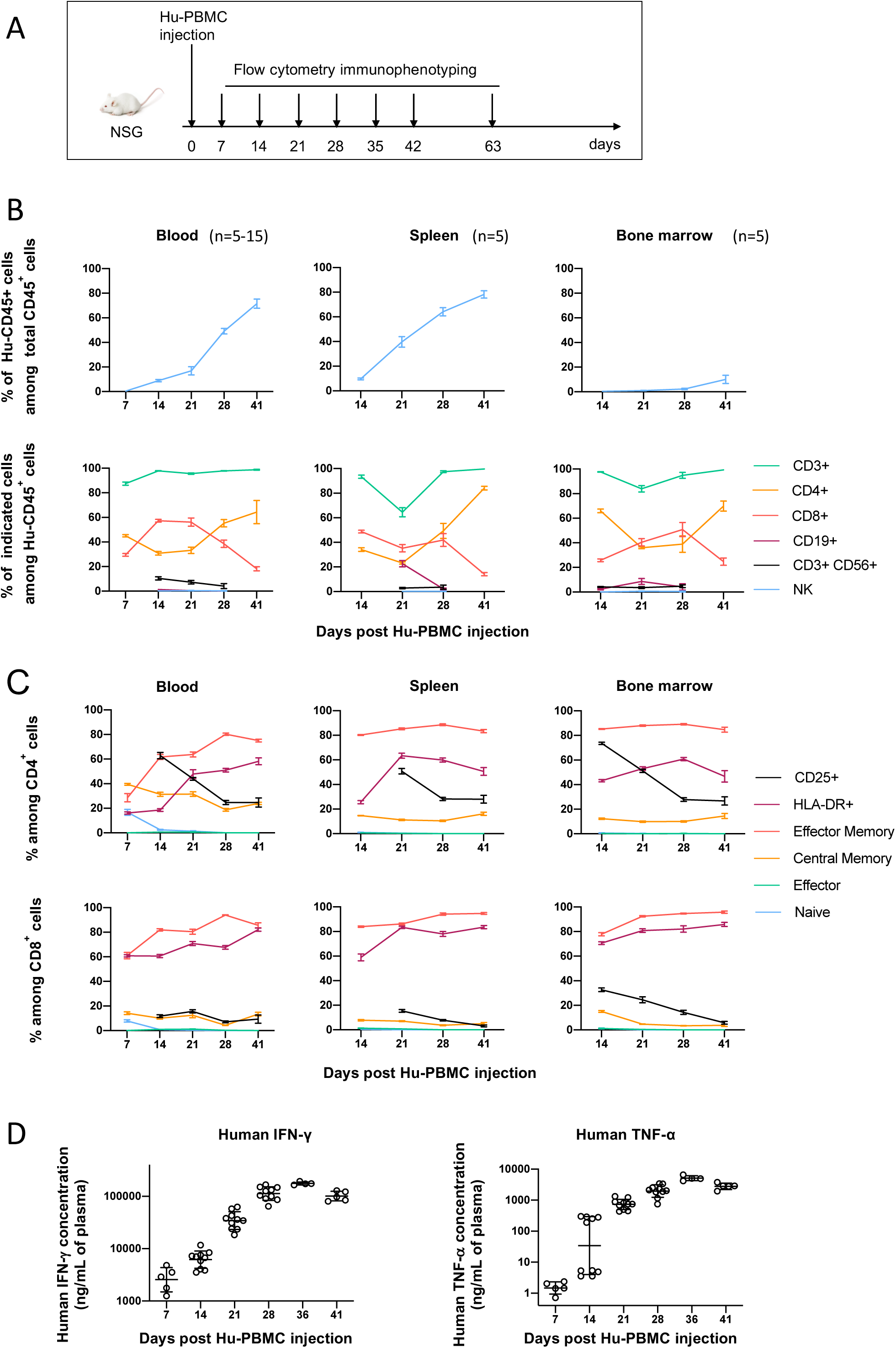
Human immune cell reconstitution upon Hu-PBMCs injection in NSG mice. **A.** Experimental design. Non-irradiated NSG mice from 8 to 14 weeks of age were injected with 10×10^6^ of Hu-PBMCs. Blood, spleen and bone marrow were recovered at illustrated time points to quantify human reconstitution by flow cytometry analysis. **B.** Changes in Hu-CD45^+^ cells reconstitution in blood (n=5-10), spleen (n=5) and bone marrow (n=5). Frequencies (%) of Hu-CD45^+^ cells relative to total CD45^+^ cells (Hu-CD45^+^ + m-CD45^+^) (upper panel) and of human immune cells subpopulations (CD3^+^ T cells, CD3^+^ CD4^+^ T cells, CD3^+^ CD8^+^ T cells, CD56^+^ CD3^+^ cells, CD19^+^ B cells and NK cells) relative to Hu-CD45^+^ cells (lower panel) at different time points. The gating strategy is described in Supplementary Fig. 1A. Data is expressed as mean ± SD of representative results obtained with one out of five donors. **C.** Changes in differentiation and activation state of CD4^+^ (upper panel) and CD8^+^ T cells (lower panel) in blood (n=5-15), spleen (n=5) and bone marrow (n=5) of reconstituted NSG mice. Frequencies (%) of effector memory, central memory, effector and naïve cells as well as activation marker-expressing cells (CD25 and HLA-DR) relative to CD4^+^ or CD8^+^ T cells were determined by flow cytometry analysis at different time points. The gating strategy is described in Supplementary Fig. 2C. Data is expressed as mean ± SD from one representative experiment out of five. **D.** From this representative experiment are shown the plasma concentrations (individual dots and geometric mean multiplied or divided by geometric SD) of human IFN-γ and TNF-α in Hu-PBMC-NSG mice at different time points. p<0,05 for both cytokines until day 28. p values were not represented for clarity and were obtained with One-Way ANOVA on log data and are adjusted with Bonferroni Holm’s correction for multiplicity.

Our results indicate that: i) A minimum dose of 5 × 10⁶ CD3+ T cell-containing PBMCs is required for consistent engraftment. ii) The maximal level of reconstitution (% Hu-CD45+ in blood) is positively correlated with the PBMC dose. iii) GvHD incidence is positively correlated with the number of PBMCs present in the inoculum. iv) The timing of GvHD onset correlates with PBMC dose, with clinical symptoms appearing earlier in mice receiving higher PBMC numbers. And v) GvHD symptoms typically emerge when Hu-PBMC levels in blood exceed 30%, though lower thresholds were occasionally observed.

Thus, we determined that injecting 5-10 × 10⁶ CD3+ T cell-containing PBMCs provided the best balance between immune reconstitution and manageable GvHD onset (**Fig. S2A-B**) and these doses were used in subsequent experiments. To account for inter-donor variability, at least at least two independent PBMC donors were included per group.

Kinetic analysis of immune reconstitution (**Fig. 2B**) revealed that Hu-CD45+ cells appeared in circulation as early as day 7–14 post-injection, reaching peak levels (∼70% in blood, ∼80% in spleen, ∼10% in bone marrow) by day 41. CD3+ T cells comprised ∼90% of engrafted Hu-CD45+ cells by day 7. Initially, CD4+ and CD8+ T cells were present in equal proportions, but from day 28 onward, CD4+ T cells became dominant. A similar trend was observed in the spleen and in bone marrow. From day 7 to day 41 post-PBMCs injection, a minor proportion of CD3+ CD56+ cells were found in the blood and spleen, and few B cells were found in the bone marrow (Fig. 2B).

We analyzed the dynamics of T cell differentiation and activation in Hu-PBMC-NSG mice, comparing them to PBMCs from healthy donors (**Fig. 2C, Fig. S2C-D**). T cells rapidly acquired an activated phenotype, with HLA-DR expression increasing from day 7 onward. CD4+ central memory T cells declined over time, while effector memory cells expanded, and CD25 expression peaked at day 14. CD8+ T cells showed earlier activation, with ∼60% becoming effector memory by day 7, expressing HLA-DR, and increasing further over time. Central memory and CD25+ cells represented 15% of CD8+ T cells and were stable from day 14. Naïve and terminal effector CD4+ and CD8+ T cells were barely detectable beyond day 14. Similar trends were observed in the spleen and bone marrow. These changes correlated with rising levels of IFN-γ and TNF-α in blood, highlighting sustained immune activation and pro-inflammatory cytokine production (**Fig. 2D**).

Collectively, our data highlight the rapid T-cell engraftment in Hu-PBMC-NSG mice but also confirm the inevitability of GvHD onset. Nevertheless, by optimizing PBMC dosing, we identified a 2–4-week therapeutic window during which T-cell-targeting immunotherapies can be evaluated without overt GvHD.

### 3- Pre-clinical evaluation of anti-PD1 in Hu-CD34^+^ mice engrafted with patient-derived xenografts (PDXs) or with tumor cell lines (cell line-derived xenograft)

Hu-CD34+ humanized mice offer several advantages over Hu-PBMC mice, including their superior reconstitution of multiple immune cell types, reduced T-cell activation, and minimal GvHD. These characteristics enable extended immunotherapy studies that are not feasible in the Hu-PBMC model. Here, we tested the ability of Hu-CD34+ mice to mount anti-tumor immune responses against PDXs or tumor cell lines following immune checkpoint inhibition (ICI) therapy.

A mini-PDX trial was conducted by injecting seven PDL-1 high small-cell lung carcinoma (SCLC) PDXs in NSG mice humanized with Hu-CD34+ cells derived from 5 different donors (**Fig. 3A and Fig. S3**). We observed that in four out of five cohorts (cohorts 1, 2, 3, and 5), anti-PD1 led to tumor-growth inhibition in at least one PDX model tested. However, in cohort 4, no response to anti-PD1 was detected, suggesting that immunotherapy outcomes in the Hu-CD34+ model can be significantly influenced by the HSC donor, independent of tumor-intrinsic factors.

**Figure 3:**
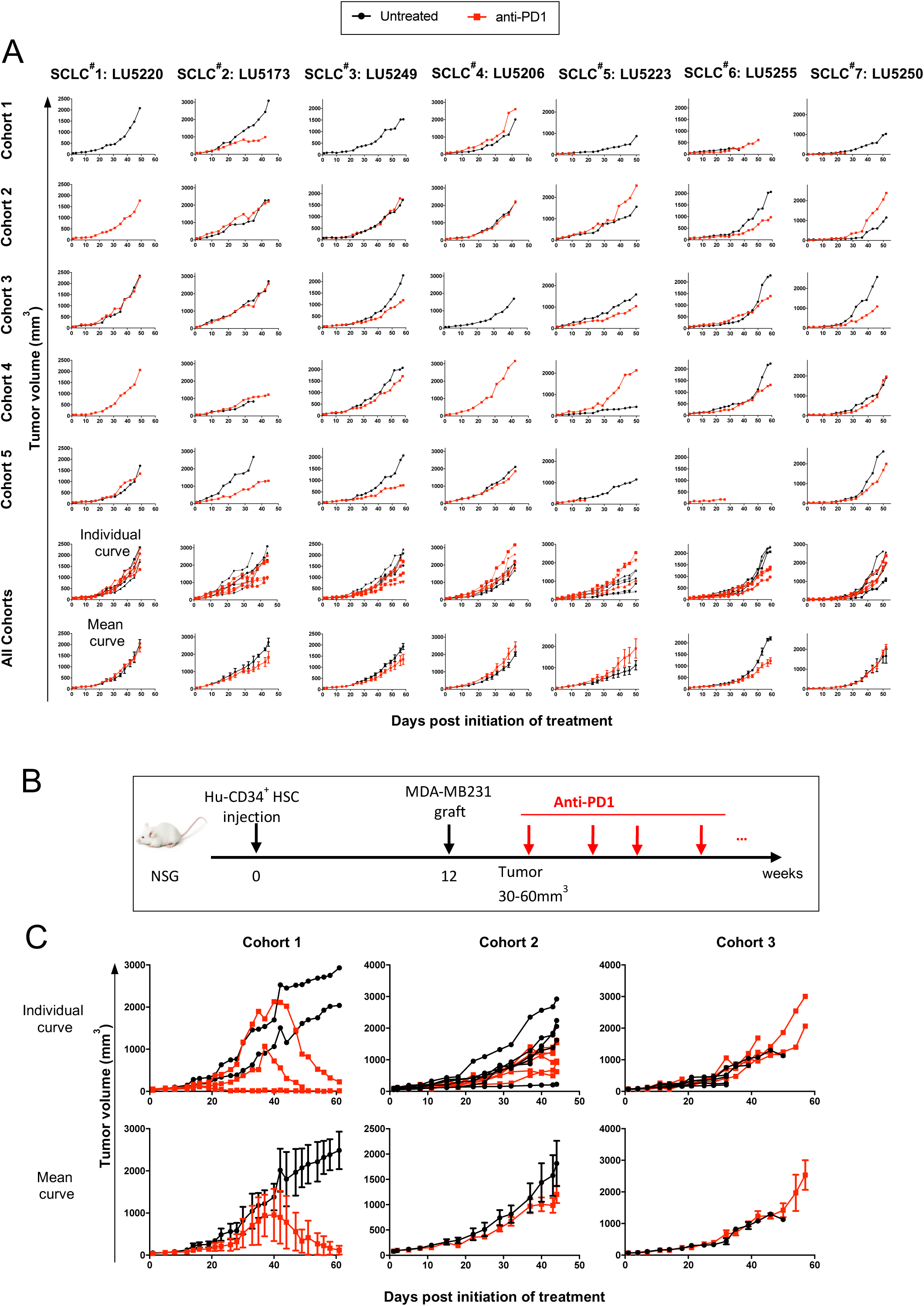
Effect of anti-PD1 mAb treatment on PDXs or cell line tumor growth in a Hu-CD34^+^-NSG mice. **A.** A N of 1 mini-PDX trial study was performed in NSG mice which were inoculated with 100×10^3^ Hu-CD34^+^ HSCs from 5 different donors. Mice reconstituted with > 25% Hu-CD45^+^ cells in PB by 12 weeks post-HSC inoculation are used. The 5 Hu-CD34^+^ cohorts were engrafted with 7 models of SCLC PDXs and mice were treated with either isotype control antibody (named “untreated”, black lines) or anti-PD1 mAb, Pembrolizumab (red lines). Tumor growth of individual mice are shown and then all cohorts are pooled as individual and mean ± SEM curves**. B.** Experimental design. Mice reconstituted with Hu-CD34^+^ cells from 3 different donors (cohort 1 to 3) were engrafted with MBA-MB231 breast tumor cell line and then treated with either isotype control antibody (black lines) or anti-PD1 mAb, Pembrolizumab (red lines). **C.** Tumor growth of individual mice (upper panel) and mean ± SEM (lower panel) are shown for the 3 separated cohorts.

To further evaluate the tumor-intrinsic sensitivity to anti-PD1, we pooled the tumor-growth curves for each PDX across all cohorts. As shown in the two bottom rows of **Fig. 3A**, SCLC PDXs #1, 4, 5, and 7 displayed no response to anti-PD1, whereas PDXs #2, 3, and 6 exhibited only weak partial responses. However, given the overall lack of strong responses and the variability in tumor growth kinetics across cohorts, it remains challenging to draw definitive conclusions regarding the tumor-intrinsic sensitivity to anti-PD1. The mild responses observed in certain PDXs may reflect variability in the dispersion of response curves rather than a consistent tumor-intrinsic susceptibility to therapy.

Similar response heterogeneity was observed when using a CDX, the MDA-MB-231 breast tumor cell line (**Fig. 3B-C**). While one cohort demonstrated complete responses in all three mice, the other two cohorts exhibited weak to no tumor regression following anti-PD1.

Collectively, these findings highlight the significant impact of the HSC donor on ICI therapy outcomes and reveal key limitations of the Hu-CD34+ model for preclinical immunotherapy evaluation.

### 4- Graft versus tumor (GvT) and GvHD in the Hu-PBMC model: a crosstalk between human tumors and immune cells

We next evaluated the impact of timing and sequence of Hu-PBMC and tumor xenograft injections (**Fig. 4A**). As shown in **Fig. 4B-C**, when PBMCs were injected 15 days prior to tumor engraftment (PDX-LCF29 or MDA-MB-231 cell line) tumor growth was completely abrogated. In contrast, in tumor-only control mice, both models grew as expected. Importantly, no overt GvHD was observed during the first four weeks (**Fig. 4D**), and all mice exhibited successful immune reconstitution, as indicated by the presence of circulating Hu-CD45+ cells in the blood (**Fig. 4E**). These findings underscore the potent GvT effect exerted by pre-engrafted PBMCs, which effectively prevent tumor growth.

**Figure 4:**
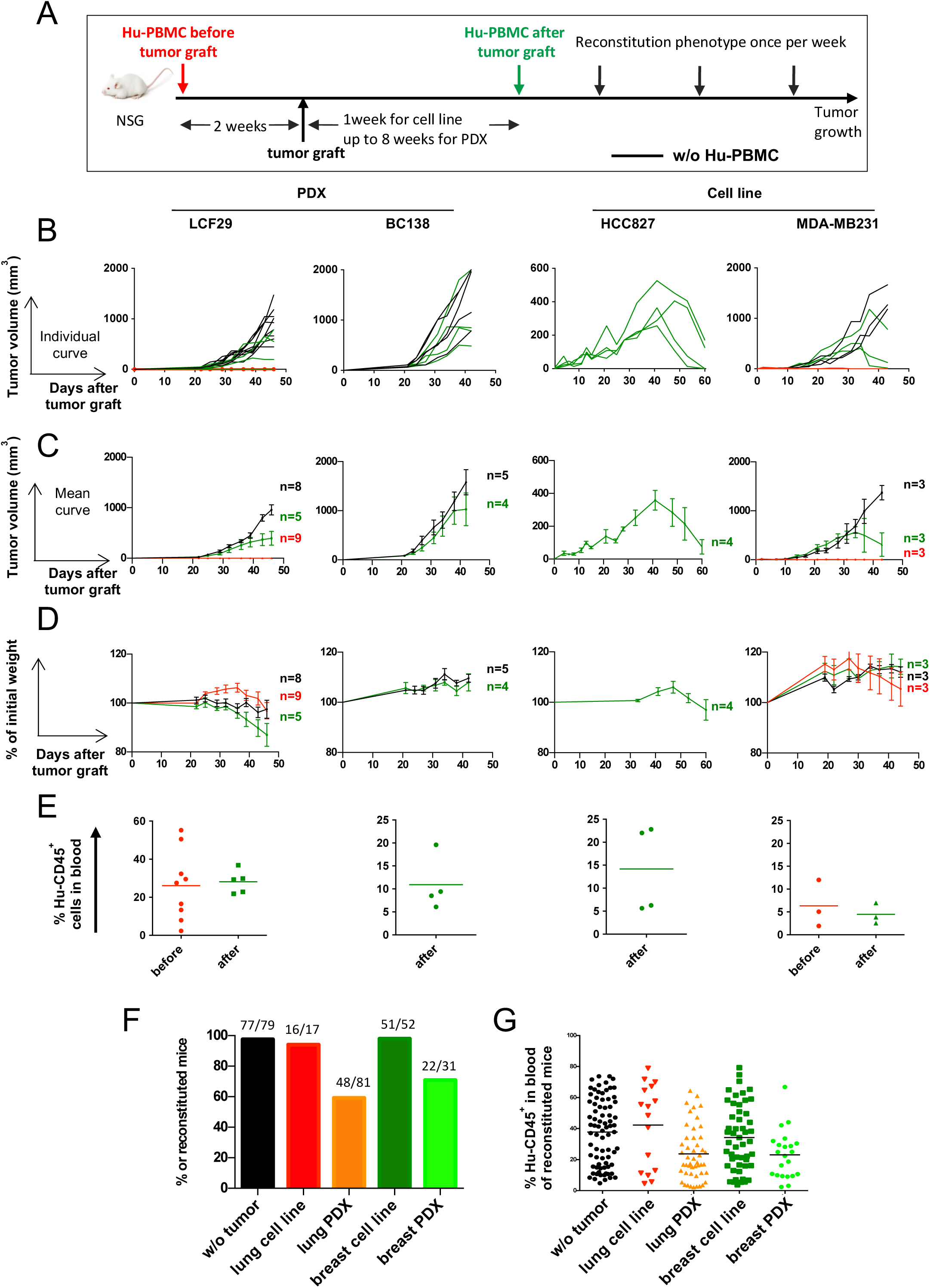
A crosstalk between human tumors and human immune cells. **A.** Experimental design. 5 or 10×10^6^ CD3^+^ T cell-containing Hu-PBMCs were injected two weeks before tumor graft (red lines) or after tumor graft was detectable (green lines); 1 week for tumor cell lines, up to 8 weeks for PDXs. As reference, tumor growth was measured in mice non injected with PBMCs (black lines). **B.** Individual tumor growth kinetics of LCF29 (NSCLC PDX), BC138 (TNBC PDX), HCC827 (NSCLC cell line) and MDA-MB231 (TNBC cell line). **C.** Tumor growth kinetics represented as mean ± SEM of individual curves shown in (B). **D.** GvHD development was followed by weight loss, represented by percentage (%) of initial weight. **C, D.** Data are represented as mean ± SEM of n=3-9 mice per group. **E.** Blood was recovered to measure Hu-CD45^+^ cells 30 days after Hu-PBMCs injection for LCF29, 22 days for BC138, 27 days for HCC827, and 20 days for MDA-MB231. **F, G.** NSG mice were not grafted with tumor (black, n=79) or grafted with NSCLC lung tumor cell line (HCC827, red, n=17), NSCLC PDX (orange, n=81), breast tumor cell line (MDA-MB231, dark green, n=52) or breast PDX (light green, n=31). Then mice were injected with 5×10^6^ to 7.5×10^6^ CD3^+^ T cell-containing Hu-PBMCs. Mice were evaluated for Hu-CD45^+^ staining in blood. **F.** Proportion of reconstituted mice. Mice were considered reconstituted when more than 2% of Hu-CD45^+^ were detected at two different time points. Numbers above bars indicates number of reconstituted mice/total PBMCs injected mice. **G.** Frequency (%) of Hu-CD45^+^ cells in reconstituted tumor-bearing mice at the maximal percentage of Hu-CD45^+^ cells detected in F (between day 30 and day 40 after tumor graft) with the mean shown for each group. Data is from a pool of several experiments.

Conversely, when PBMCs were injected after tumors became palpable, all four tested tumor models continued to grow. Compared to tumor-only controls, three of the tested tumors (PDX-LCF29, PDX-BC138, and MDA-MB-231) exhibited slower growth, suggesting a delayed but detectable GvT effect. For tumors with slower intrinsic growth kinetics (such as HCC827 and MDA-MB-231), prolonged monitoring revealed eventual tumor regression, further supporting a delayed GvT effect in these models. For the more aggressive PDX-BC138 model, at 4 weeks post-engraftment up to 10% of the tumor infiltrate consisted of Hu-CD45+ immune cells, primarily CD4+ T cells, and low proportions of CD8+ T cells and NK cells (**Fig. S4**).

Circulating Hu-CD45+ cell levels, kinetics and intensity of GvHD, varied across tumor models and did not predict tumor growth control (Fig. 4D-E). The mechanisms underlying GvHD onset in this model may involve the mismatch of major histocompatibility complex (MHC) molecules between the engrafted human immune cells and the mouse host tissue (xeno-GvHD), and the allo-response of T cells from the donor attacking different human MHC molecules present in the tumor (13,17). When comparing pooled data from multiple experiments, we observed lower frequencies of Hu-PBMC reconstitution in PDX than in CDX models in or tumor-free controls, suggesting that PDXs negatively impact PBMC engraftment (**Fig. 4F**). Additionally, the maximum PBMC reconstitution levels, varied across conditions (**Fig. 4G**). Despite high inter-mouse variability in Hu-CD45+ percentages, tumor-free and CDX-engrafted mice exhibited superior PBMC reconstitution compared to PDX-bearing mice, indicating that tumor type can influence immune cell dynamics.

Overall, these findings highlight a bidirectional interplay between engrafted human immune cells and tumors. While PBMCs can mediate a robust GvT effect against certain tumors, PDXs can, in turn, hinder immune cell reconstitution. Consequently, to standardize the Hu-PBMC tumor model for subsequent studies, we established a protocol wherein tumors were always engrafted prior to PBMC injection, with concurrent monitoring of immune reconstitution and mouse body weight to account for the above-mentioned variables.

### 5- Pre-clinical evaluation of anti-PD1 in the Hu-PBMC model

To assess the suitability of the Hu-PBMC model for immunotherapy studies, we evaluated the response to anti-PD1 of Hu-PBMC NSG mice engrafted with eight different NSCLC-PDXs (**Fig. 5A**). Four PDX models (LCF13, LCF15, LCF16, and LCF25) were completely resistant to anti-PD1, while three (LCF17, LCF22, and LCF29) exhibited mild responses. Only one model (LCF26) demonstrated significant sensitivity to PD1 blockade (**Fig. 5B).**

**Figure 5:**
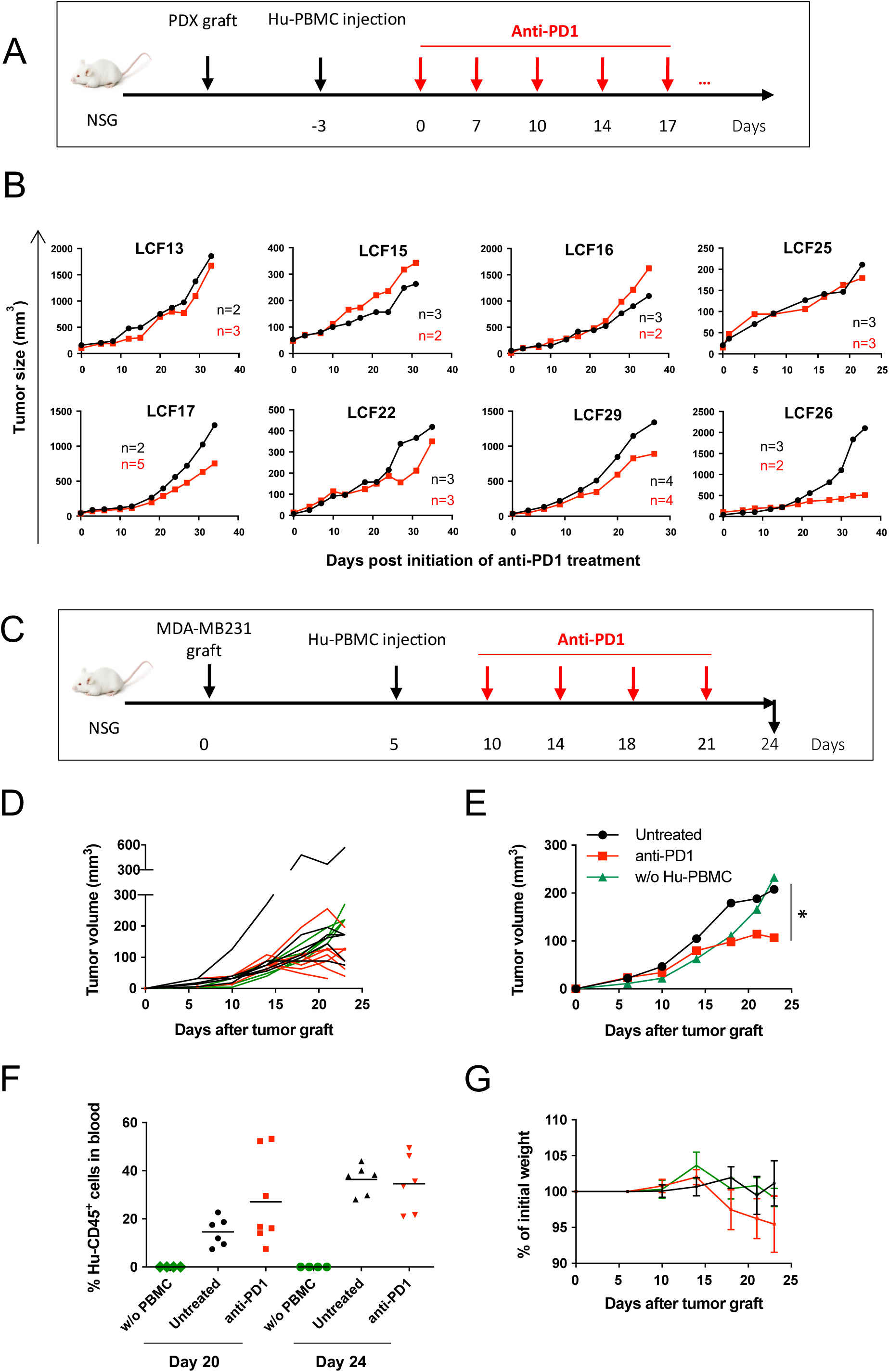
Effect of anti-PD1 treatment on tumor growth in Hu-PBMC-NSG model. **A.** Experimental design. NSCLC PDX-bearing NSG mice were injected with 5 to 7.5×10^6^ CD3^+^ T cell-containing Hu-PBMCs and treated 3 days after Hu-PBMCs injection either with anti-PD1 (Nivolumab) twice per week (red line, n=2-5) or with PBS (black line, n=2-4). Mice showing > 2% Hu-CD45+ cells in circulation at least at two different time points during the experiment were included. **B.** Tumor growth kinetics with 8 NSCLC PDX models. Data is represented as mean of the tumor size **C.** Experimental design. NSG mice were grafted with MDA-MB231 breast tumor cells. Once tumor was detectable (tumor size range= 10-30 mm^3^), mice were injected with 5×10^6^ CD3^+^ T cell-containing Hu-PBMCs. Five days after Hu-PBMCs injection, mice received PBS (black, n=5) or anti-PD1 mAb (Nivolumab) (red, n=7) twice per week. Control mice did not receive Hu-PBMCs (green, n=4). **D.** Tumor growth kinetic of one representative experiment out of two, shown as individual curves. **E.** Tumor growth kinetic shown as the mean from mice shown in the figure 5D. *global p value <0.05 was calculated for the comparison between untreated and anti-PD1 treated group and obtained with a Two-Way ANOVA-Type. **F**. Frequency (%) of Hu-CD45^+^ cells in blood at day 20 and day 24. **G.** GvHD development was followed by weight loss. Data is represented as mean ± SEM.

Immune reconstitution varied significantly across PDX-bearing mice, and anti-PD1 correlated with reduced PBMC engraftment in 6 of the 8 models tested (**Fig. S5A).** Using the MDA-MB-231 model, we observed a modest delay in tumor growth following anti-PD1 (**Fig. 5C-E**). At day 20 post-PBMC injection, anti-PD1-treated mice exhibited a slight increase in circulating Hu-CD45+ cells (**Fig. 5F**), coinciding with the onset of weight loss (**Fig. 5G**). Notably, weight loss in anti-PD1-treated mice may reflect treatment-related toxicity rather than purely anti-tumor effects. At the experimental endpoint, both untreated and anti-PD1-treated tumors exhibited high levels of Hu-CD45+ immune cell infiltration (**Fig. S5B**), with no discernible differences between treatment groups.

Although tumor control by anti-PD1 was mild and heterogeneous, similar findings were independently reproduced in a separate laboratory (**Fig. S5C-F**), demonstrating the reproducibility of these results. However, the observed responses underscore the limited efficacy of anti-PD1 in Hu-PBMC models, likely due to their unique immune environment.

### 6- Pre-clinical evaluation of other anti-tumor T-cell directed therapies in the Hu-PBMC model

Given the observed limited efficacy of anti-PD1 and the heterogeneous responses across tumor models, we explored other immunotherapies that may better harness T-cell-mediated tumor rejection. We focused on two additional promising T-cell-directed therapies: an IL-2-based immunotherapy (IL-2Cx)(18) and a T-cell engager (EpCAM-CD3 bispecific antibody(19), which enhance anti-tumor immunity by promoting the activation of specific T-cell subsets and offer alternative mechanisms that could overcome limitations observed with anti-PD-1.

We first evaluated the IL-2Cx which preferentially activates NK and CD8+ T cells and has demonstrated efficacy in models of transplantable tumors (14,20). In the LCF29 PDX-bearing Hu-PBMC model, anti-PD1 had no significant impact on tumor growth, whereas IL-2Cx treatment induced marked tumor inhibition (**Fig. 6A-C**) and accelerated GvHD (**Fig. 6D**). Flow cytometry revealed that IL-2Cx increased the proportion of CD4+ relative to CD8+ T cells in circulation (**Fig. S6**). IL-2Cx-expanded CD4+ T cells displayed an effector phenotype and did not express FoxP3 (data not shown), indicating that IL-2Cx primarily activated conventional (non-Treg) CD4+ T cells. At 40 days post-PDX engraftment, tumors from IL-2Cx-treated mice exhibited a mild increase in Hu-CD45+ immune cell infiltration, dominated by CD3+ T cells, with a shift toward effector CD4+ T-cell enrichment (**Fig. 6E).**

**Figure 6:**
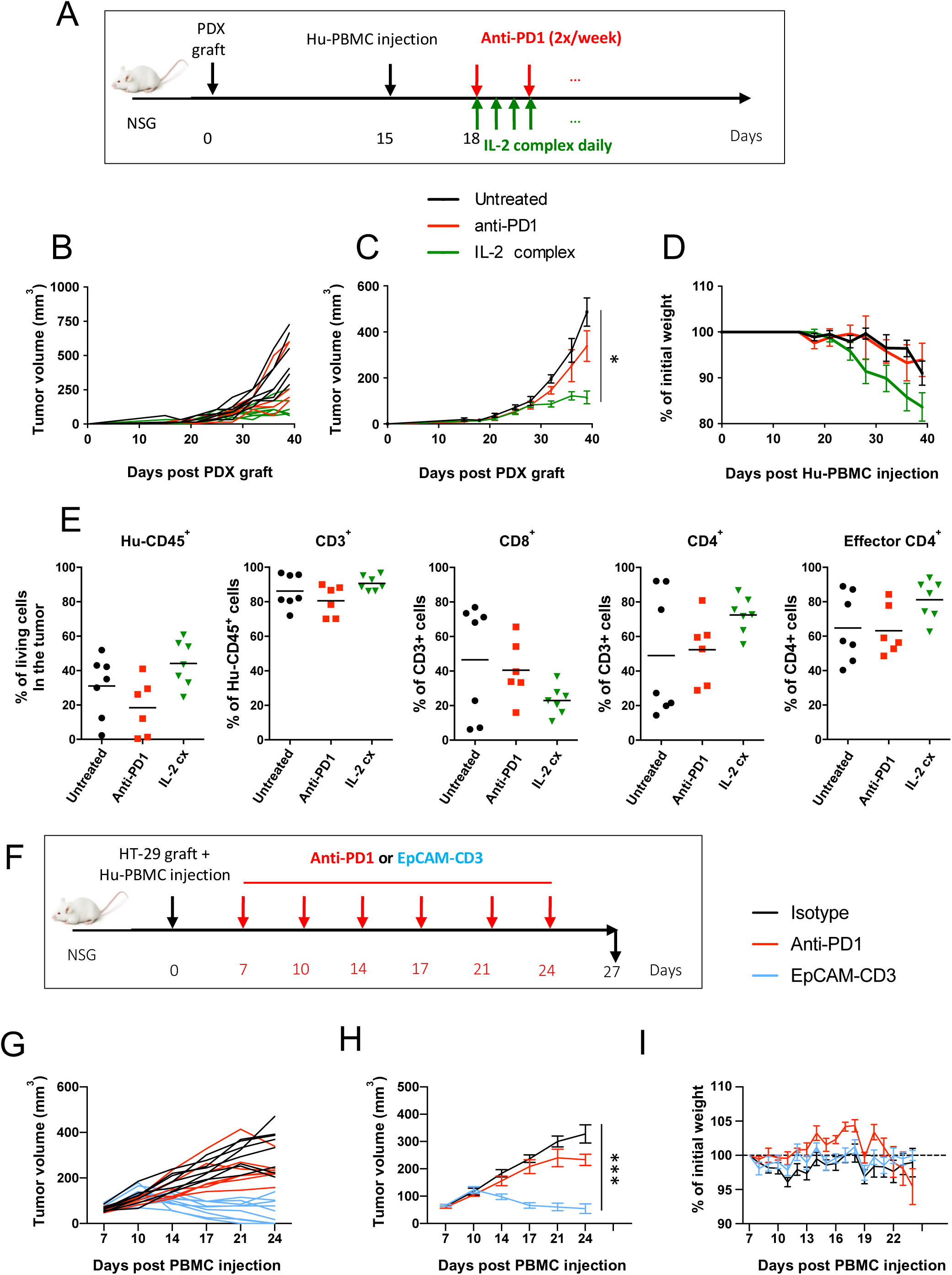
T-cell driven therapies show high anti-tumor efficacy in the Hu-PBMC model. **A.** Experimental design. NSCLC LCF29 PDX-bearing NSG mice were injected with 7.5×10^6^ CD3^+^ T cell-containing Hu-PBMCs. Mice were treated either with PBS (black, n=7), anti-PD1 mAb (Nivolumab) (red, n=6), or IL-2Cx (green, n=7). Only reconstituted mice were included (i.e., mice showing > 2% Hu-CD45^+^ cells in circulation in at least two different time points during the experiment). **B and C.** Tumor growth kinetics represented as individual curves (**B**) and as mean ± SEM of n=6-7 mice per group (**C**). *global p value was <0.05 and obtained with a Two-Way ANOVA-Type with Bonferroni-Holm’s correction for multiplicity. **D.** GvHD onset was followed by weight loss represented as mean ± SEM. **E.** Flow cytometry analysis of tumor infiltrating cells was done 40 days after PDX engraftment (25 days after Hu-PBMCs injection). Frequencies (%) of the indicated human immune cell populations are shown. **F.** Experimental design. NSG mice were injected on day 0 with HT-29 tumor cells and with 10×10^6^ Hu-PBMCs. On day 7, when tumor reached 45-125 mm^3^, mice were treated with anti-PD1 mAb (Nivolumab), or with isotype control or with an EpCAM-CD3. **G.** Individual HT-29 tumor growth kinetics. **H.** Tumor growth kinetics represented as a mean ± SEM of n=7-9 mice per group**. I.** GvHD development was followed by weight loss. Data are represented as mean ± SEM of n=7-9 mice per group. ***global p value was <0.0001 and was obtained with a Two-Way ANOVA-Type with Bonferroni-Holm’s correction for multiplicity.

Next, we tested the EpCAM-CD3 bispecific antibody, which engages T cells through TCR activation (anti-CD3 Ab arm) and targets tumors via EpCAM recognition on tumor surface. A anti-PD1 induced a slight tumor growth delay, whereas EpCAM-CD3 led to significant tumor regression, with complete tumor resolution observed in 3 out of 9 mice (**Fig. 6F-H**), underlying the potential of bispecific antibodies in inducing potent anti-tumor responses.

Overall, these results highlight the utility of the Hu-PBMC model for evaluating immunotherapies, particularly those that directly and robustly activate T cells. Unlike PD-1 blockade, which shows heterogeneous responses likely influenced by donor variability, therapies such as IL-2Cx and EpCAM-CD3 bispecific antibodies induce strong T-cell activation, leading to more consistent and reproducible immune responses in this model.

## Discussion

Humanized mouse models enable in vivo study of human immune responses and immunotherapies, but systematic comparisons remain scarce (5). Direct side-by-side evaluation of Hu-PBMC and Hu-CD34+ mice, CDX and PDX, using standardized criteria has been lacking. By assessing immune composition, donor variability, tumor responsiveness, and therapeutic outcomes, our study provides a framework to clarify strengths and limitations for future applications.

Consistent with previous studies (21), Hu-PBMC and Hu-CD34+ HSC injections yield distinct immune profiles. Hu-PBMC rapidly establishes a T cell–dominated environment with activated CD3+ T effectors (22), while Hu-CD34+ supports multilineage (T, B, NK, myeloid) reconstitution but requires longer engraftment (12). Consistent with prior reports in Hu-CD34+ models, donor-dependent variability strongly influenced immune composition and therapy response (23). Interestingly, CDX favored PBMC reconstitution over PDX, likely due to their less immunosuppressive microenvironment and potential MHC differences.

PD-1 blockade showed limited efficacy in Hu-CD34+ models despite clinical success in NSCLC (24–26). This mirrors other reports that Hu-CD34+ models lack the immunological complexity for robust checkpoint efficacy (22). Impaired APC maturation (8,27) and absence of human cytokines may restrict T-cell activation. By contrast, our NSCLC PDX trial with anti-PD-1 showed ∼50% responses, close to clinical rates (28), underscoring donor and tumor-specific influences. Thus, while Hu-CD34+ models capture some aspects of clinical heterogeneity, future models require improved myeloid and stromal interactions.

In contrast to the limited effects observed with PD1-blockade, Hu-PBMC models responded strongly to T-cell engaging therapies. IL-2Cx induced CD4+ T cell–driven antitumor activity, differing from CD8+/NK-driven effects in syngeneic mice (18), but consistent with humanized settings (29,30). EpCAM-CD3 bispecifics also induced tumor regression, including complete responses, validating their relevance for T-cell engagers evaluation (31,32). Hu-PBMC models’ ability to capture T-cell activation dynamics makes them valuable for evaluating therapies that bypass classical antigen presentation and rely on direct T-cell engagement.

Despite their value, both models require refinement. Hu-CD34+ offers longer study windows and multilineage reconstitution but is costly and lack key human cytokines. Transgenic strains expressing human cytokines (e.g., IL-15, GM-CSF, IL-3, CSF) may enhance immune function and better model human immune-tumor interactions (5,33).

The immune-avatar strategy, co-engrafting early-passage PDXs with autologous immune cells could enhance personalization and avoid mismatching (30,34–36). Given the difficulty of obtaining Hu-CD34+ cells from cancer patients, Hu-PBMC models offer a feasible alternative for implementing this strategy. However, challenges remain, particularly GvHD and limited study duration. The generation of MHC I/II knockout models (37,38) may help address these limitations.

Humanized mouse models remaino key for CAR-T evaluation, enabling monitoring of expansion, antitumor activity, donor-specific responses, and toxicities -such as cytokine release syndrome or off-target effects-before clinical trials (39,40).

## Conclusion

While humanized mice are indispensable for preclinical immuno-oncology, refinements are needed to better model human immune complexity. Emerging transgenic models and immune-avatar strategies hold promise for bridging preclinical and clinical studies, enabling more precise and effective cancer immunotherapies.

## Acknowledgements

We thank the mouse facility and flow cytometry platforms from Institut Curie and Animal Platform CRP2-UMS 3612 CNRS-US25 Inserm-IRD from Faculty of Pharmacy, Paris Descartes University. We thank the Pharmacy of Institut Curie for providing us with Nivolumab. We thank Nathalie Amzallag for project management and Laurent Andrieu for feedback on this manuscript. We would also like to thank Ercole Rao and Christian Biel (Sanofi Biologics, Frankfurt) for providing the EpCAM-CD3 bispecific Ab.

