## Supplementary figures for "A comprehensive analysis of humanized mouse models for the study of cancer immunotherapies"

### Supplementary Figure 1

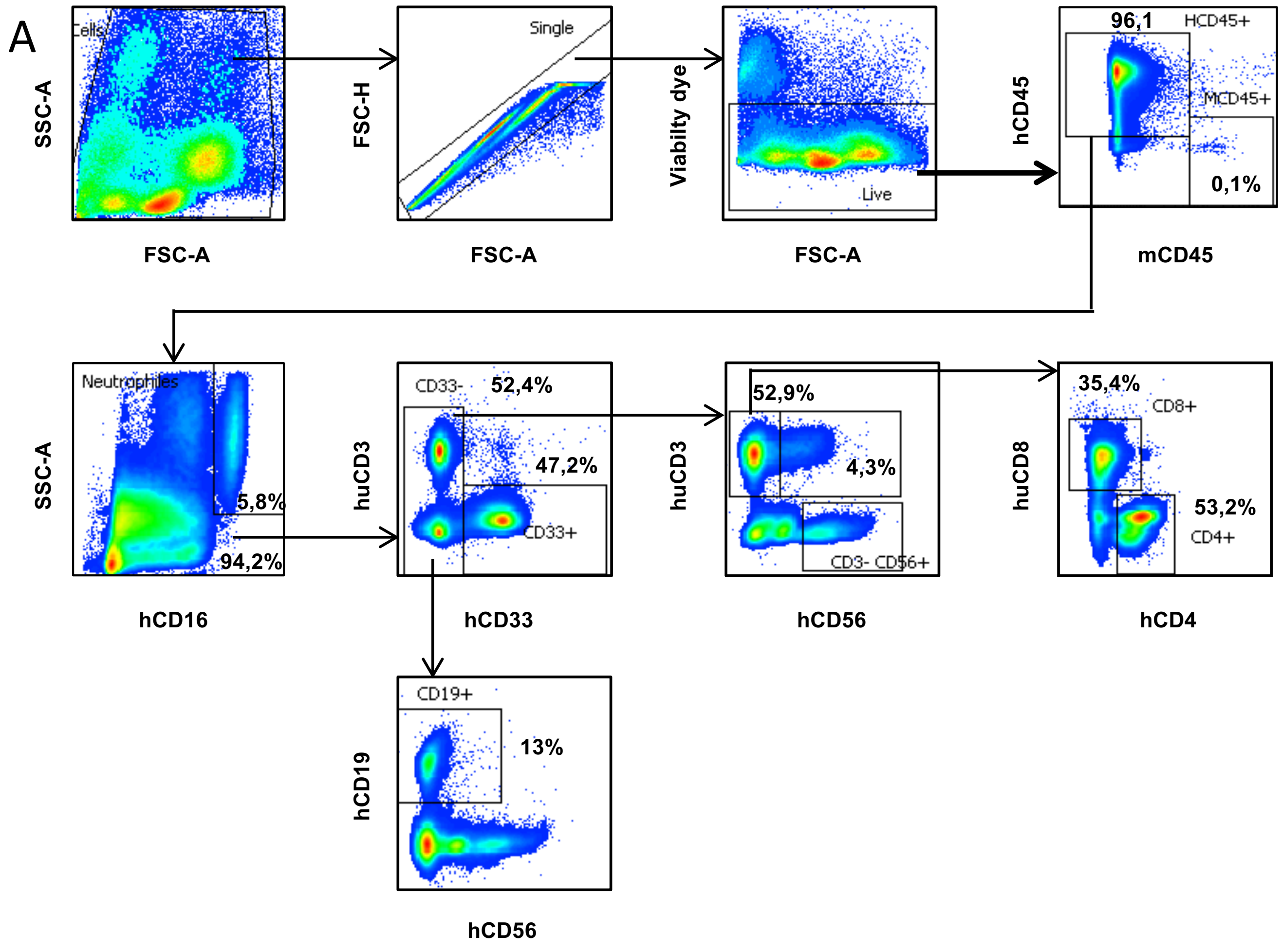

### Supplementary Figure 1

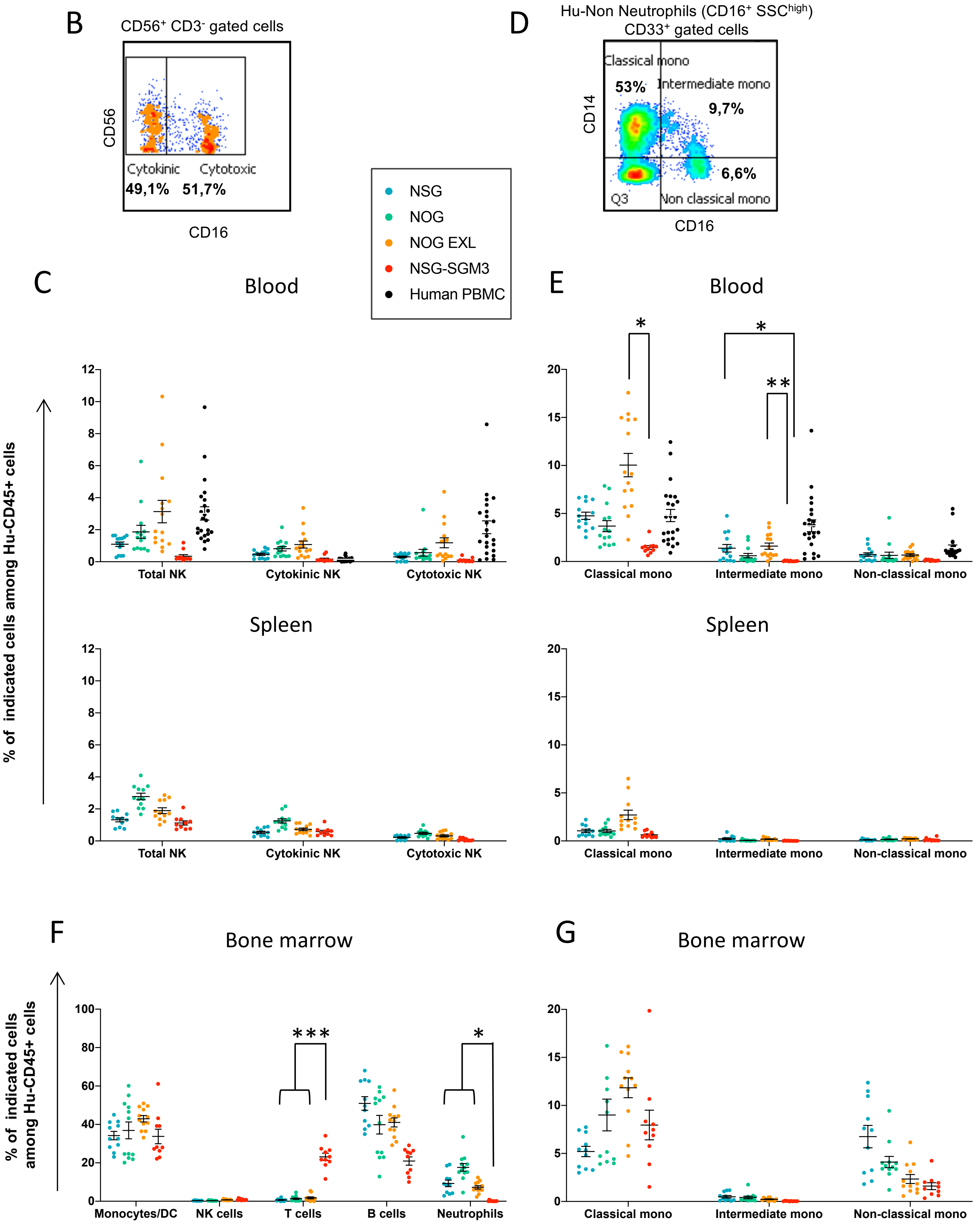

**Supplementary Figure 1: Characterization of the NK and Myeloid cell reconstitution upon Hu-CD34$^{+}$ HSC injection. A.** Flow cytometry gating strategy for analysis of human immune cell subpopulations, shown in blood of Hu-CD34$^{+}$ HSC-humanized mouse at week 18 after humanization as in Figure 1C. Similar gating strategy was used for Hu-PBL-NSG mouse. **B.** Flow cytometry gating strategy used to define cytotoxic (CD16$^{+}$), and cytokinic (CD16$^{-}$) NK cells among Hu-CD56$^{+}$CD3$^{-}$ gated cells (see Supplementary Figure 1A for gating strategy) in blood from Hu-CD34$^{+}$-humanized mice at week 18. **C.** Percentage of NK cell subpopulations relative to Hu-CD45$^{+}$ cells in blood and spleen at week 18. **D.** Flow cytometry gating strategy used to define classical (CD14$^{+}$ CD16$^{-}$), intermediate (CD14$^{+}$ CD16$^{+}$), and non-classical (CD14$^{-}$ CD16$^{+}$) monocytes among Non Neutrophils (CD16$^{+}$ SSC$^{high}$) CD33$^{+}$ gated cells (see Supplementary Figure 1A for gating strategy) in blood from Hu-CD34$^{+}$-humanized mice at week 18. **E.** Percentage of monocyte subpopulations relative to Hu-CD45$^{+}$ cells in blood and spleen at week 18. **F.** Distribution of the Hu-CD45$^{+}$ cell subpopulations relative to Hu-CD45$^{+}$ cells in bone marrow of different mouse strains at week 18. **G.** Percentage of monocyte subpopulations relative to Hu-CD45$^{+}$ cells in bone marrow at week 18.

Data were obtained from mice reconstituted with 2 or 3 different Hu-CD34+ HSC donors with n=10-15 mice per mouse strain for blood, bone marrow and spleen data. Data is expressed as individual dots and mean ± SD. *p<0,05, ****p* ≤ 0.001 was obtained with a One-Way Analysis of Variance on log-transformed with Tukey's correction for multiplicity.

### Supplementary Figure 2

A

| Number of injected CD3-containing Hu-PBMC x10$^6$ (Total injected Hu-PBMCs) | Frequency of reconstituted mice | Mean % of maximum reconstitution | Frequency of GvHD | Mean day of GvHD onset (range) | Mean % (range) of reconstitution at GvHD onset |
|---|---|---|---|---|---|
| 20-30 (30-50) | 2/2 (100%) | 49,9% | 2/2 (100%) | 16 (14-18) | 32,7% (20,5-44,8) |
| 10 (16-30) | 14/15 (93%) | 42,3% | 11/14 (78%) | 34 (12-51) | 38% (15,5-66,3) |
| 5 (9-12) | 17/18 (94%) | 34,1% | 11/17 (65%) | 42,3 (29-63) | 39,2% (10,7-73,7) |
| 1-3 (2-4) | 0/3 (0%) | / | / | / | / |

B

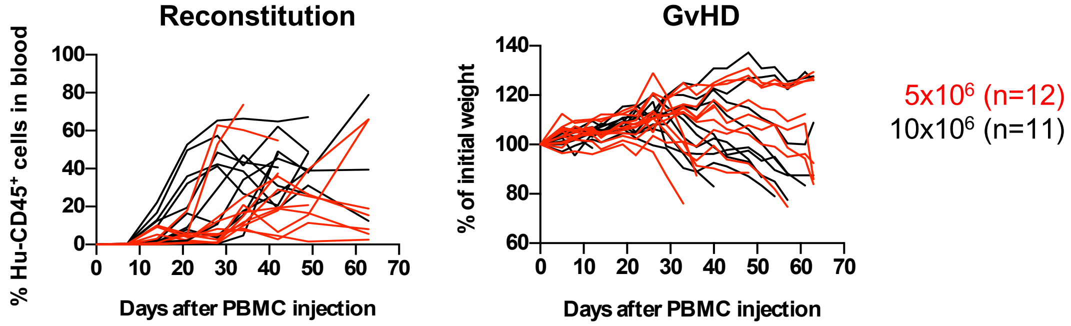


C

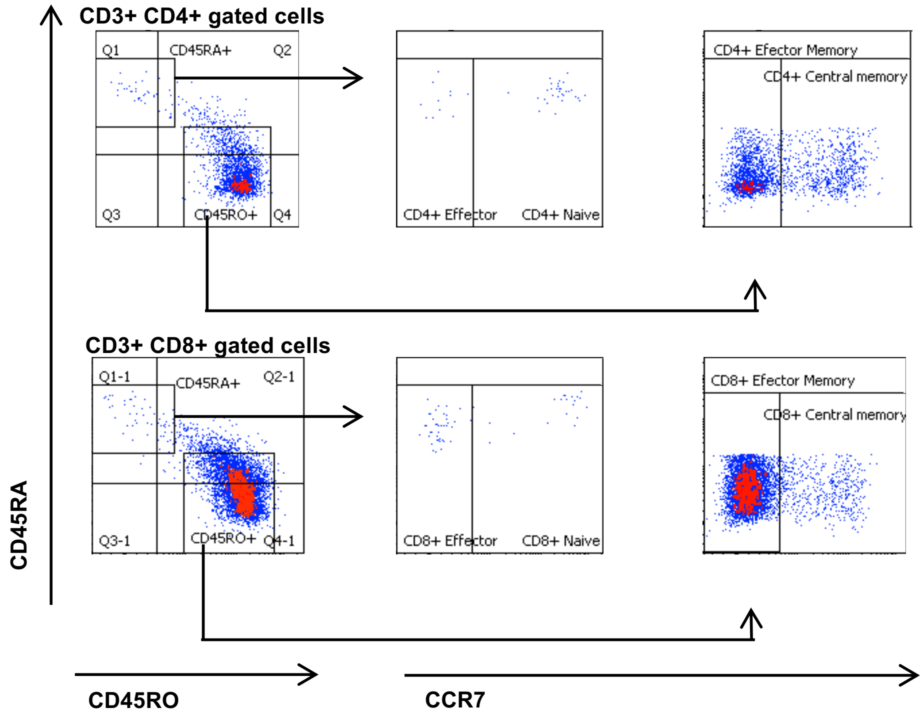


D

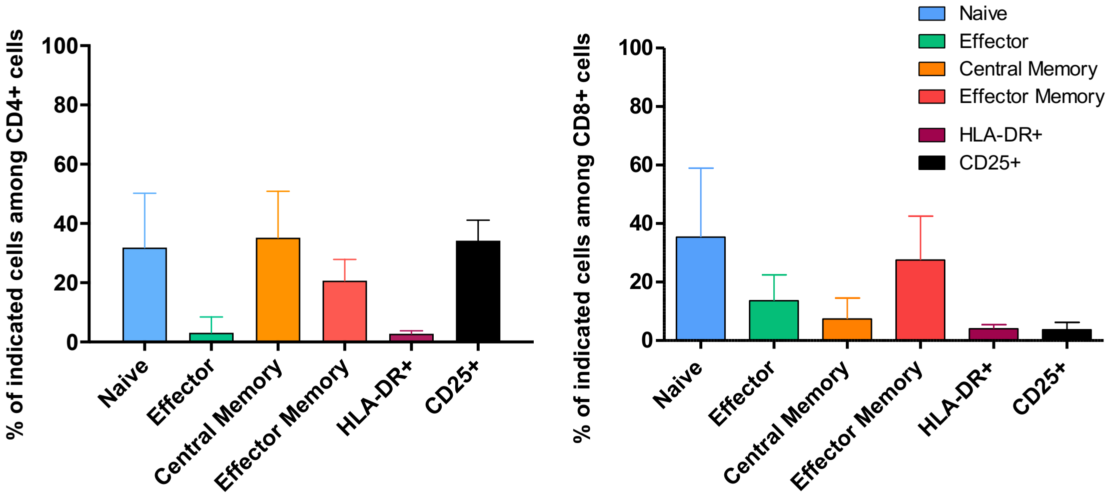

**Supplementary Figure 2 : Hu-PBMCs reconstitution and GvHD development in Hu-PBMC-NSG mice**

**A.** Non-irradiated NSG mice from 8 to 14 weeks of age were injected with different amount of Hu-PBMCs (1 to 30x$10^6$) and the Hu-PBMCs engraftment efficiency (measured by the % of Hu-CD45+ cells in blood) and GvHD development (measured by the weight loss) are shown in the table. **B.** NSG mice injected with 5x$10^6$ (red lines, n=12) or 10x$10^6$ (black lines, n=11) of $CD3^+$ T cell-containing Hu-PBMCs were monitored for reconstitution by quantification of blood circulating Hu-$CD45^+$ (left panel) cells and bodyweight (right panel). Results shown are percentage of initial weight. Statistical analysis on hCD45+ cells was calculated until day 48 with a Two-Way ANOVA on log data for reconstitution and on raw data for GvHD and were not different between the 2 groups. **C.** Flow cytometry gating strategy used to define naïve ($CD45RA^+$ $CCR7^+$), central memory ($CD45RA^-$ $CCR7^+$), effector memory ($CD45RA^-$ $CCR7^-$), and effector cells ($CD45RA^+$ $CCR7^-$) among Hu-$CD4^+$ or Hu-$CD8^+$ gated cells in blood from Hu-PBMC-humanized mice at day 14 post-humanization. **D.** T cell differentiation/activation state in blood from healthy donor. Frequencies (%) of naïve, memory, effector memory, and effector cells as well as activation marker-expressing cells (CD25 and HLA-DR) relative to $CD4^+$ or $CD8^+$ T cells of PBMC from healthy donors (n=3-4) used to inject in NSG mice, determined by flow cytometry analysis. Data is expressed as mean ± SD.

### Supplementary Figure 3

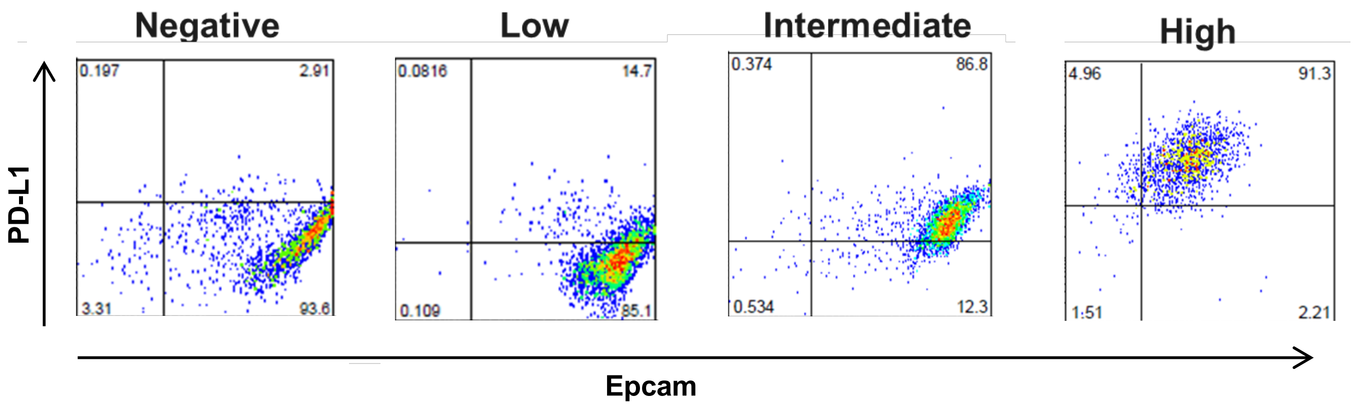


**Supplementary Figure 3: PD-L1 expression by SCLC PDXs.**
30 SCLC PDX tumors were screened by FACS for their PD-L1 expression level and examples of low / intermediate or high expression are illustrated. 7 high PD-L1 expressers were identified and used in the mini-PDX trial study presented in Figure 3A.

### Supplementary Figure 4

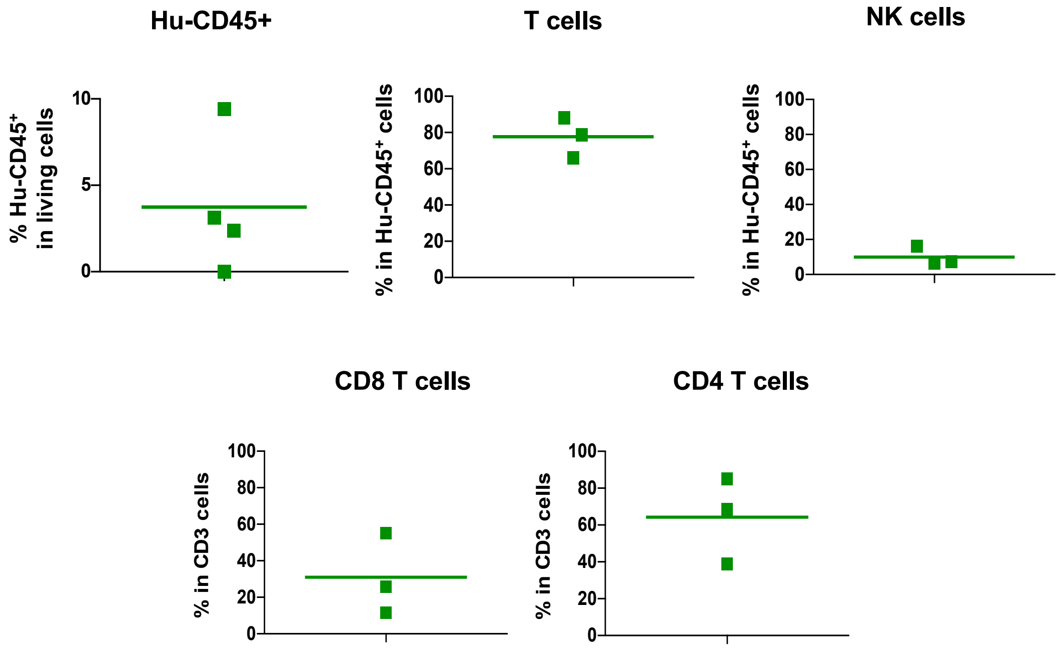


**Supplementary Figure 4: Human immune infiltration in TNBC PDX**

TNBC PDX (BC138) was sampled 28 days after PBMC injection. Tumor infiltration of human immune cells (Hu-CD45+) was assessed by flow cytometry. Frequencies (%) of tumor infiltrating CD56 (NK cells), CD3 (T cells), CD8 (T cytotoxic cells) and CD4 (T helper cells) are individually shown.

### Supplementary Figure 5

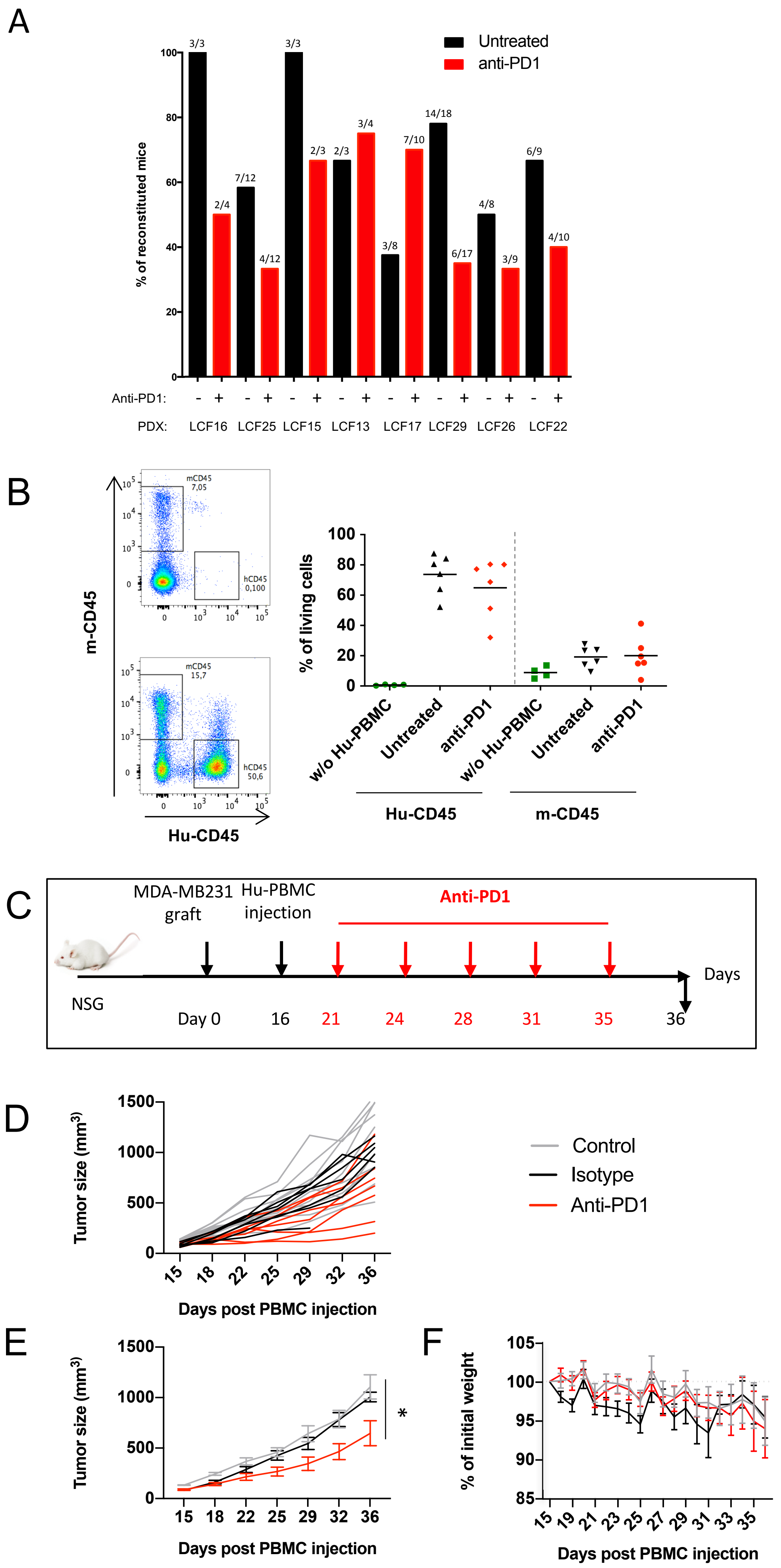

**Supplementary Figure 5: Effect of anti-PD1 treatment on lung PDXs or on MDA-MB231 tumor growth in a Hu-PBMC-NSG model.**

**A.** Details of the proportion of reconstituted tumor-bearing mice shown in Figure 5A/B, in 8 NSCLC PDX bearing mice models treated or not with anti-PD1. Mice were considered reconstituted when more than 2% of Hu-CD45$^+$ were detected in blood. Numbers indicated the reconstituted mice/total Hu-PBMCs injected mice. **B.** Mice from the experiment described in Figure 5C were analyzed for the content in Hu-CD45+ or m-CD45+ cells at the tumor site. Representative dot plot showing the frequency (%) of tumor infiltrating m-CD45$^+$ and Hu-CD45$^+$ cells in non-PBMC injected mouse (upper panel, left) and in PBMC-injected mouse (lower panel, left). Quantification of m-CD45$^+$ and Hu-CD45$^+$ cells in tumors from the 3 different groups (without injected PBMC, receiving PBMC and non-treated or anti-PD1 treated) is shown on the right panel. **C.** Scheme of the experimental design. NSG mice were injected with MDA-MB231 breast tumor cells and on day 16 mice were injected ip with $10x10^6$ Hu-PBMC. On day 21, when tumor reached 61-144 $mm^3$, mice were treated bi-weekly with 10 mg/Kg of anti-PD1 (Nivolumab) or 10mL/kg of Isotype anti-HEL huIgG4 and the control group was not treated. **D**. Individual MDA-MB231 tumor growth kinetics and **E** Tumor growth kinetics represented as a mean ± SEM of the individual curves shown in (D). *global p value <0.05 was calculated for the comparison between Isotype Control and anti-PD1 treated group and obtained with a Two-Way ANOVA-Type. **F**. GvHD development was followed by weight loss, represented by percentage (%) of initial weight when treatments started. Data are represented as mean ± SD of n=6-10 mice per group.

### Supplementary Figure 6

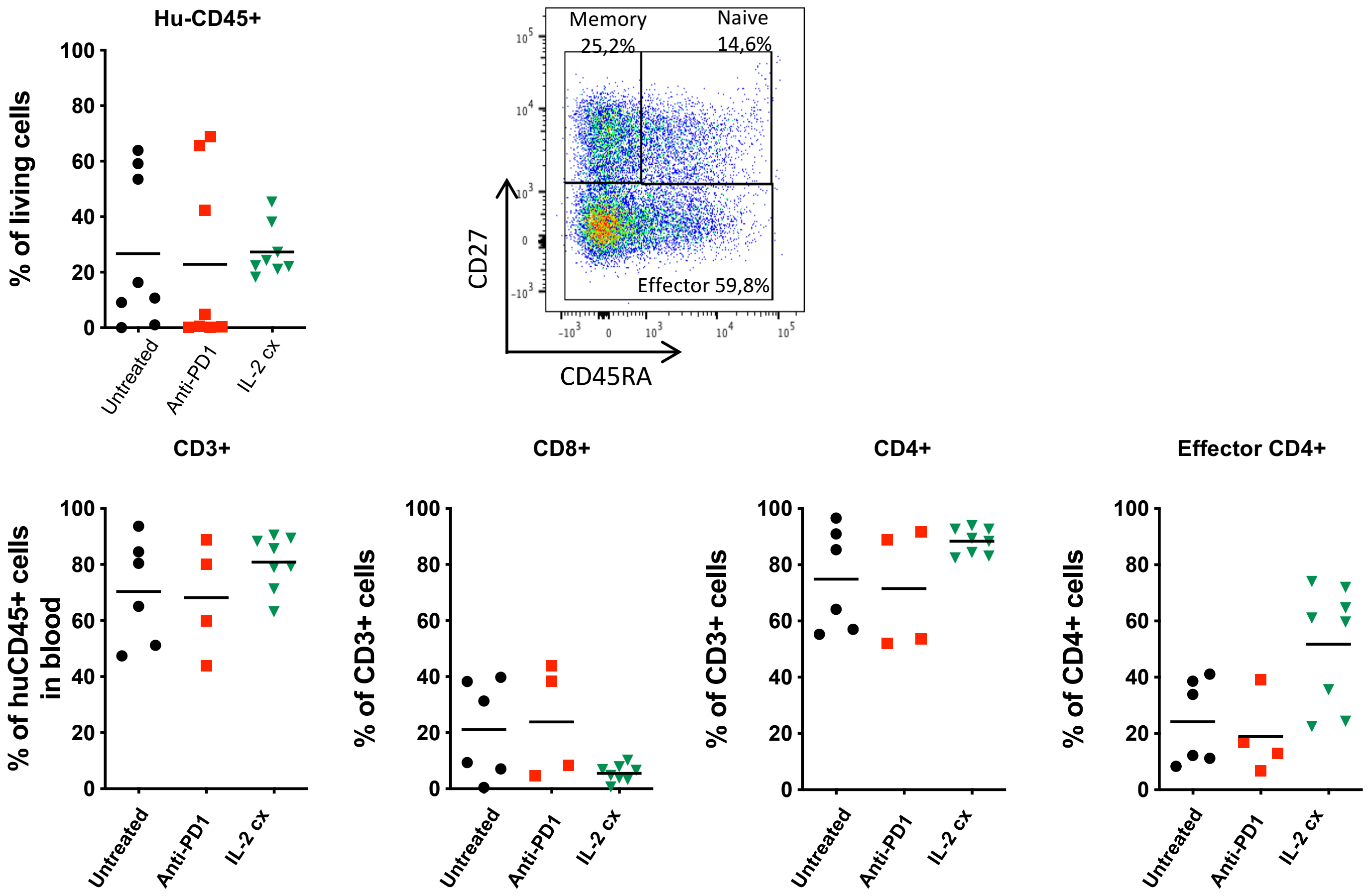


**Supplementary Figure 6: IL-2Cx effect on human immune cell reconstitution in the blood of NSCLC-PDX LCF29 Hu-PBMC NSG mice.**
LCF29 PDX-bearing NSG mice were treated as described in Figure 6A. The percentages of the indicated immune cell populations in the blood, analyzed at day 21 after Hu- PBMCs injection, are shown. Insert illustrates gating strategy to define effector CD4 T cells.
